# TopLib: Building and searching top-down mass spectral libraries for proteoform identification

**DOI:** 10.1101/2024.11.12.623220

**Authors:** Kun Li, Haixu Tang, Xiaowen Liu

## Abstract

Mass spectral library search is a widely used approach for spectral identification in mass spectrometry (MS)-based proteomics. While numerous methods exist for building and searching bottom-up mass spectral libraries, there is a lack of software tools for top-down mass spectral libraries. To fill the gap, we introduce TopLib, the first software package designed for building and searching top-down spectral libraries. TopLib utilizes an efficient spectral representation technique to reduce database size and improve query speed and performance. We systematically evaluated various spectral representation techniques and scoring functions for top-down spectral clustering and search. Our results demonstrated that TopLib is 140 times faster and achieves better reproducibility in proteoform identification compared with conventional database search methods in top-down MS.

## 1. Introduction

Mass spectrometry (MS) is a powerful technique for identifying and quantifying peptides, proteins, and proteoforms in complex biological samples [1]. In a typical tandem mass spectrometry (MS/MS)-based proteomics experiment, two types of MS spectra are generated: MS1 spectra measure the molecular masses of peptides or proteoforms and MS/MS spectra measure the mass-to-charge ratios (*m*/*z*) and intensities of fragments of peptides or proteoforms [2]. There are two main approaches for identifying MS/MS spectra: database search and spectral library search [3-6]. While database search methods compare query spectra against a protein/proteoform sequence database for spectral identification [3, 4], spectral library search methods compare query spectra against the spectra of peptides or proteoforms collected in a pre-built spectral library for spectral identification [5, 6]. Compared with database search, spectral library search leverages the intensity information of fragment ions observed in experimental spectra, enhancing the sensitivity in spectral identification [7].

Bottom-up and top-down MS are two commonly used methods in MS-based proteomics [8]. Bottom-up MS analyzes peptides generated through enzymatic digestion of proteins, while top-down MS directly examines intact proteoforms [9]. In recent years, top-down MS has become the preferred method for identifying and characterizing intact proteoforms [10], but there is a lack of software tools designed for building and searching top-down spectral libraries for proteoform identification. Although numerous approaches have been developed for building and searching spectral libraries in bottom-up MS [6, 11-13], these methods cannot be directly applied to top-down MS due to the differences between the mass spectra generated in top-down and bottom-up MS. For example, top-down mass spectra typically contain more high charge state ions than bottom-up spectra, and spectral deconvolution [14, 15], which converts a complex mass spectrum into a list of monoisotopic masses, is commonly required as a preprocessing step in top-down mass spectral analysis.

Efficient representation of mass spectra is critical for building and searching spectral libraries [12]. While exploiting all fragment ions in MS/MS spectra enhances the sensitivity in spectral identification, it demands extensive storage space and slows down spectral library search. Efficiently representing mass spectral can greatly reduce the size of spectral libraries and accelerate spectral library search, without substantially compromising the sensitivity of spectral identification. Common approaches for representing spectra in bottom-up spectral libraries include bin-based methods [11], in which a mass spectrum is divided into *m*/*z* bins and each *m*/*z* bin is represented by its signal intensity, and deep learning models [12], which convert mass spectra into vector representations.

In spectral library-based methods, similarity scoring functions for ranking spectrum-spectrum-matches (SSMs) are essential for improving the accuracy of spectral clustering and the sensitivity of spectral identification. In bottom-up MS, various similarity or distance functions have been used for evaluating SSMs, including spectral dot product [16, 17], Pearson correlation coefficient [18], Spearman’s rank correlation coefficient [19], and Euclidean distance [12]. Additionally, a relative entropy-based distance function has been applied to spectral library searches in MS-based metabolomics [20].

We introduce TopLib, the first software package designed for building and searching top-down spectral libraries. We conducted a systematic assessment of various spectral representation methods, similarity and distance functions, and spectral clustering algorithms for building and searching top-down spectral libraries. Experiments on top-down MS data generated from colorectal cancer cells demonstrated that spectral library search using TopLib identified many spectra missed by database search, increased search speed by 140 times, and improved the reproducibility of proteoform identification compared with database search.

## 2. Results

### 2.1 Overview of TopLib

TopLib consists of four functions for building top-down mass spectral libraries (Fig. 1). The first function is top-down spectral deconvolution [14, 15] (Fig. 1a), which simplifies complex mass spectra by converting isotopic peaks of precursor and fragment ions into monoisotopic neutral masses. The second function identifies proteoform-spectrum-matches (PrSMs) by searching deconvoluted mass spectra against a protein sequence database [4, 21, 22] (Fig. 2b). TopFD [14] and TopPIC [4] are employed for spectral deconvolution and database search in TopLib. The third function builds a top-down comprehensive spectral library using deconvoluted top-down mass spectra, along with associated meta data and proteoform identification results stored in text files. The fourth function groups mass spectra in the comprehensive library into clusters and generate representative mass spectra for spectral clusters.

**Fig. 1:**
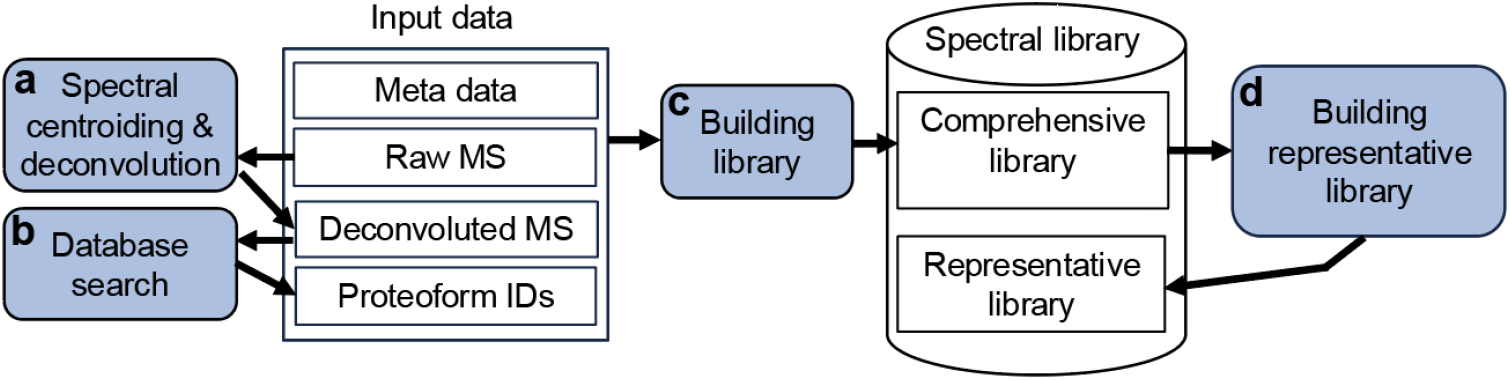
Overview of TopLib. Four functions are used for building top-down spectral libraries: (a) spectral centroiding and deconvolution, which convert mass spectra into monoisotopic neutral mass lists; (b) database searching of deconvoluted mass spectra for proteoform identification; (c) building a comprehensive library; and (d) generating a representative library from the comprehensive library. The representative library contains a representative spectrum for each cluster generated from the comprehensive library.

**Fig. 2:**
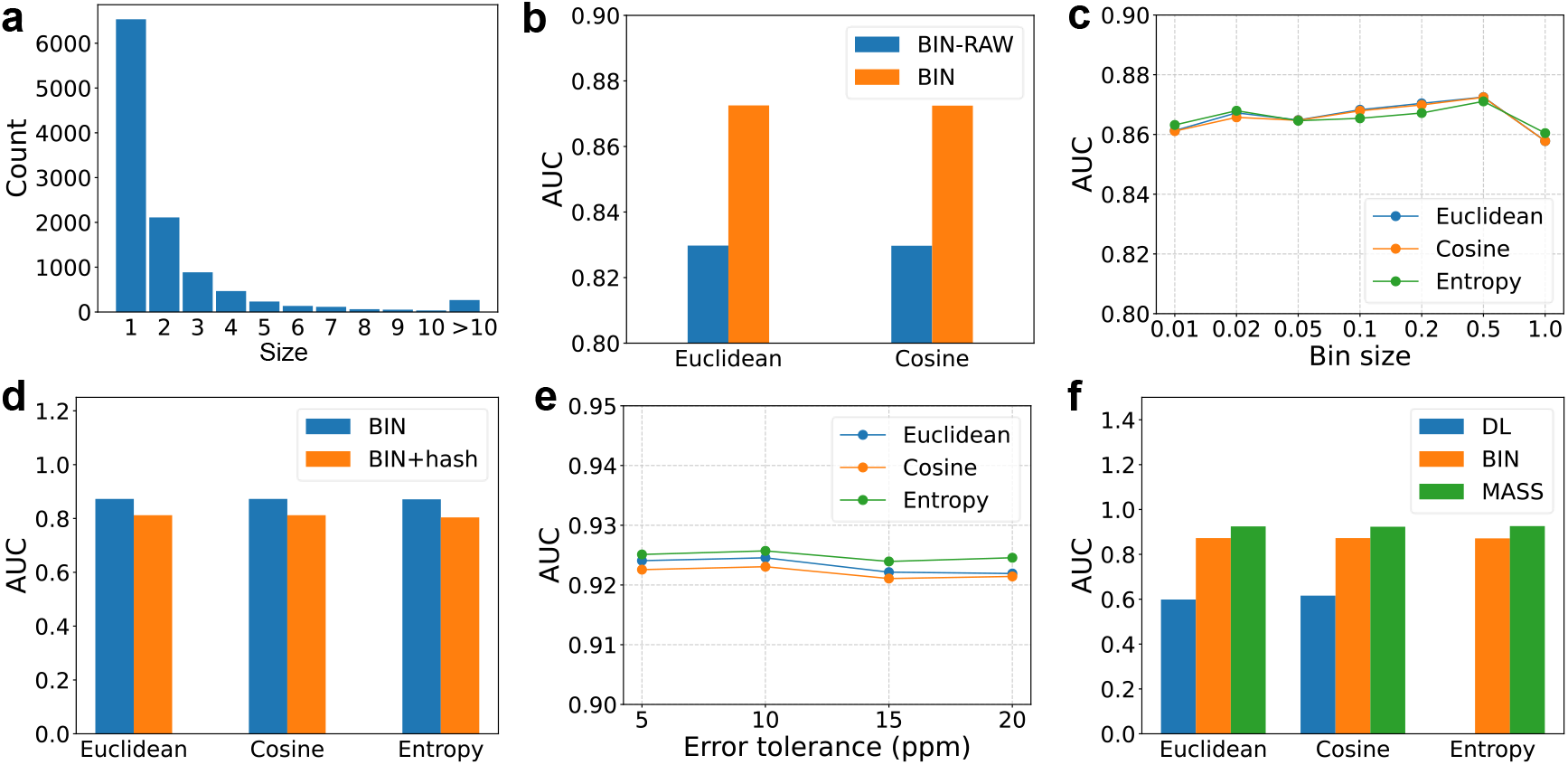
Evaluation of the DL, BIN, and MASS representation methods and Euclidean distance, cosine distance, and the entropy-based distance for top-down mass spectra. The 50 highest intensity deconvoluted fragment masses in each spectrum are used for spectral representation. (a) Distribution of the sizes of the 10,893 groups reported from the SW480-3D data set. (b) Comparison of the BIN-RAW and BIN representations on the SW480-SPE data set using a bin size of 0.5. (c) Comparison of various bin sizes in the BIN representation on the SW480-SPE data set. (d) Comparison of the BIN representation with and without a hashing function on the SW480-SPE data set using a bin size of 0.5. (e) Comparison of various settings for the error tolerance in the MASS representation on the SW480-SPE data set. (f) Comparison of the three spectral representation methods on the SW480-SPE data set with a bin size of 0.5 for the BIN representation and an error tolerance of 10 ppm for the MASS representation. The entropy-based distance is not used for the DL representation because mass intensity information is not available in the representation.

### 2.2 Evaluation data sets

We generated evaluation data sets using a top-down MS data set described in McCool *et al*. [23], in which 3-dimensional (3D) separation coupled with top-down MS was utilized to analyze proteins extracted from SW480 cells. Technical triplicates were included in the data set. The first replicate of the data set, referred to as the SW480-3D data set, was used to obtain two evaluation mass spectral data sets. The SW480-3D data set contained 54 MS data files and 75,605 MS/MS spectra, which were centroided, deconvoluted, and searched against the UniProt human proteome sequence database for proteoform identification (see Methods). A total of 37,566 PrSMs were identified with a 1% spectrum-level false discovery rate (FDR). After removing possible duplicated proteoform groups and inconsistent PrSMs (see Methods), 28,913 PrSMs remained. The PrSMs were divided into 10,893 groups based on their proteoform identifications and charge states. The average size of the groups was 2.65 (Fig. 2a). Then we removed all groups with size 1, and the remaining 4,359 groups with 22,379 spectra are referred to as the SW480-3D spectral groups.

The first evaluation set was generated to assess similarity or distance functions for SSMs. We randomly sampled 5,000 spectrum pairs from same groups and 5,000 spectrum pairs from different groups from the SW480-3D spectral groups. For each different-group spectrum pair, the precursor mass difference was restricted to the range of [0, 200] Dalton (Da). Cosine similarity, calculated using a bin-based spectral representation with a bin size of 0.5 (see Section 3.4 in Methods), was employed to compare these two types of spectrum pairs. As expected, the cosine similarity scores were significantly higher for the same-group pairs compared with the different-group pairs (Supplemental Fig. S1a). To make the evaluation task harder, we reduced the distinguishability between these two types of spectrum pairs by shifting some deconvoluted fragment masses in the same-group spectra (see Methods) to lower their similarity scores (Supplemental Fig. S1b). The evaluation set, containing the 5,000 same-group pairs with shifted masses and the 5,000 different-group spectrum pairs, is referred to as the SW480 spectral pair evaluation (SW480-SPE) data set.

The second evaluation set was designed to assess the performance of spectral clustering methods. Since many SW480-3D spectral groups have distinct precursor masses, one group can easily be separated from others based solely on their precursor masses. To test the accuracy of spectral clustering when two or more groups share similar precursor masses, we changed the precursor masses of some spectra in the SW480-3D spectral groups, ensuring that most spectra could not be correctly clustered using precursor masses alone (see Methods). The resulting spectral set is referred to as the SW480 group paired spectrum evaluation (SW480-GPSE) data set.

### 2.3 Spectral representations and distance functions

We evaluated three representation methods (see Methods) for deconvoluted top-down MS/MS spectra. Since top-down MS/MS spectra often contained many isotopic peaks for each fragment ion, spectral deconvolution was employed to convert these isotopic peaks into neural monoisotopic masses to simplify the data. As a result, a deconvoluted mass spectrum can be viewed as a list of (mass, intensity) pairs or (*m*/*z*, intensity) pairs, where the *m*/*z* values correspond to the monoisotopic peaks of the deconvoluted masses. The first representation method is a deep learning-based encoding approach [12], where a deconvoluted spectrum with (*m*/*z*, intensity) pairs is converted into a vector of size 32. The second method allocates the intensities of deconvoluted masses to bins based on their *m*/*z* values [11]. The third method simplifies each spectrum by retaining only the *k* most intense (mass, intensity) pairs for representation. These methods are referred to as DL (deep learning-based), BIN (bin-based), and MASS (mass-intensity pair) representations, respectively.

We first used the SW480-SPE set to compare the performance of the BIN representation (with log transformation) and the bin-based representation without log transformation of mass intensities (see Methods), referred to as BIN-RAW. For a spectral pair with a cosine similarity *s*, the cosine distance is defined as 1-*s*. Pairwise spectral distances were computed using Euclidean distance, cosine distance, or an entropy-based score [20] (see Methods). Compared with BIN-RAW, BIN demonstrated better discriminative power in distinguishing between same-group and different-group spectral pairs using Euclidean and cosine distances (Fig. 2b). Consequently, log transformation of mass intensities was used as the default setting for computing Euclidean and cosine distances for the BIN and MASS representations. However, it was not applied to the entropy-based score, as log transformation is already integrated into the score. For the DL representation, we used the method in Bittremieux *et al*. [12] without applying intensity log transformation.

We evaluated the BIN representation with various settings for the bin size on the SW480-SPE data set. The highest area under receiver operating characteristic curve (AUC) value was achieved with a bin size of 0.5 Thomson (Th) for Euclidean distance, cosine distance, and the entropy-based score (Fig. 2c). Based on the results, the bin size 0.5 was selected as the default setting for the BIN representation. Additionally, we experimented a hash function [11] to reduce the BIN representation (the vector size is 3000 for the default *m*/*z* range [200, 1700] Th with a bin size of 0.5) to a small 800-dimensional vector. However, the hash function significantly reduced the discriminative power compared with the representation without hashing (Fig. 2d).

We also assessed various error tolerance settings for matching deconvoluted fragment masses in the MASS representation on the SW480-SPE data set. The distribution of the errors of fragment masses in the 5,000 same-group spectrum pairs (Supplemental Fig. S2) indicated that ±1 Da errors are common in deconvoluted fragment masses. Therefore, ±1 Da errors were allowed in matching fragment masses. In addition, only masses with the same charge state were matched. That is, two mass charge pairs (*m*_1_, *c*_1_) and (*m*_2_, *c*_2_) were considered to be matched if (1) *c*_1_ = *c*_2_ and (2) |*m*_1_ – *m*_2_| < *e* or 1.00235-*e* < |*m*_1_ – *m*_2_| < 1.00235 + *e*, where 1.00235 Da is an estimated average mass difference between two neighboring isotopic masses of a fragment [24]. The best AUC was obtained with an error tolerance of 10 ppm (Fig. 2e), which was chosen as the default error tolerance for the MASS representation.

We evaluated the discriminative ability of the three representation methods using the default error tolerance and various settings for the number *k* of fragment masses kept in a mass spectrum: 25, 50, 75, 100, 150, and 200 (see Methods) on the SW480-SPE data set. The highest AUC was obtained at *k* = 50 for the BIN and DL representations and at *k* = 100 for the MASS representation (Supplemental Tables S1, S2, and S3). As a result, we selected *k* = 50 as the default setting for both the BIN and DL representations. Although *k* = 100 yielded the best AUC for the MASS representation, we opted to use *k* = 50 as the default setting to reduce the spectral library size. Furthermore, experimental results for spectral clustering (see Section 2.4) indicated that *k* = 50 provided better clustering performance than *k* = 100.

Using the default settings, the MASS representation achieved the highest discriminative ability among the three methods (Fig. 2f), indicating that the DL and BIN representations may lose some information of the deconvoluted masses compared with the MASS representation, reducing their ability to accurately differentiate the same-group and the different-group spectrum pairs. For the MASS representation, no significant differences were observed across the three distance metrics.

### 2.4 Spectral clustering

We developed a Top-down spectral Clustering method, TopCluster, to group spectra in a top-down spectral library. In TopCluster, hierarchical clustering is first performed based on precursor masses, followed by hierarchical or DBSCAN [25] clustering using cosine distance to further divide spectra into clusters (see Methods). The three spectral representation methods, with their default parameter settings, were evaluated and benchmarked against spectral-cluster (version 1.1.2) [26] on the SW480-GPSE data set with 22,379 spectra from 4,359 clusters. In spectral-cluster, a probabilistic score serves as the similarity function for spectral pairs, and a greedy approach is employed for clustering.

The clusters in the SW480-GPSE data set, generated based on proteoform identifications reported by TopPIC [4] (see Section 2.2), were used as the ground truth in the evaluation. Clustering performance was assessed using three metrics: the ratio of clustered spectra, the ratio of incorrectly clustered spectra, and the completeness of clustering (see Methods). TopCluster with the BIN and MASS representations and hierarchical clustering outperformed other methods by producing a higher ratio of clustered spectra and achieving better completeness for a given ratio of incorrectly clustered spectra (Fig. 3a, 3b). TopCluster also demonstrated comparable performance with Euclidean distance and the entropy-based score (Supplemental Fig. S3) and with the DBSCAN clustering method (Fig. 3d, 3e, and Supplemental Fig. S4). Additionally, we compared the performance of the MASS representation with two settings for the parameter *k* (50 and 100) and found that *k* = 50 outperformed *k* = 100 in clustering accuracy (Supplemental Fig. S5).

**Fig. 3:**
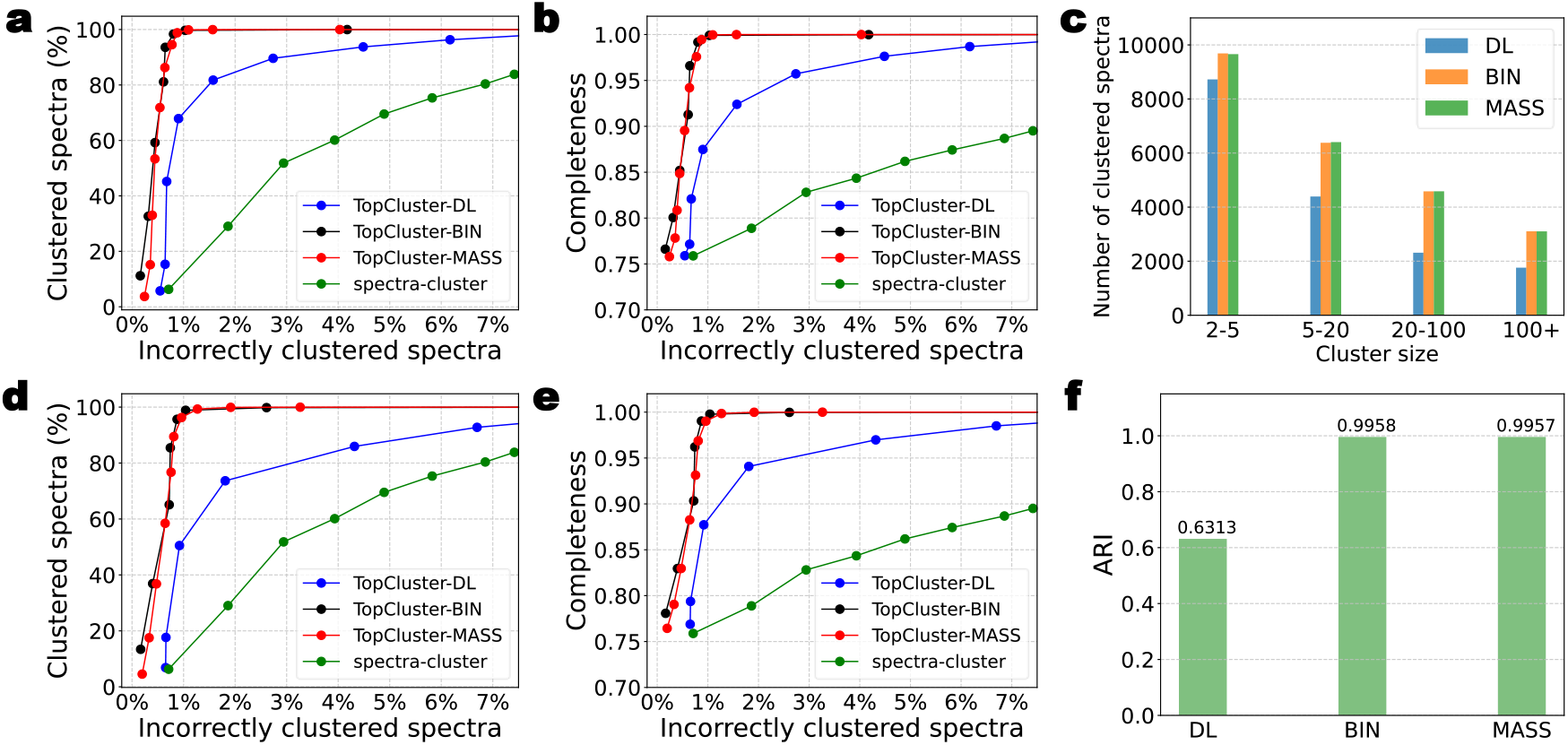
Comparison of spectral clustering accuracy of spectral-cluster and TopCluster using three spectral representation methods on the SW480-GPSE data set. (a) and (d) show the ratio of incorrectly clustered spectra against the ratio of clustered spectra for hierarchical clustering and DBSCAN, respectively. (b) and (e) plot the ratio of incorrectly clustered spectra against clustering completeness for hierarchical clustering and DBSCAN, respectively. (c) and (f) give the sizes and ARIs of the clusters reported by TopCluster using distance cutoffs of 0.161 for the DL representation, 0.796 for the BIN representation, and 0.876 for the MASS representation.

We further compared the performance of the three representation methods using a cosine distance cutoff of 0.161 for the DL representation, 0.796 for the BIN representation, and 0.876 for the MASS representation. At these cutoffs, the three methods reported similar ratios of incorrectly clustered spectra: 1.00% for DL, 1.02% for BIN, and 1.01% for MASS. TopCluster with the BIN and MASS representations identified more non-singleton clusters (BIN: 4,265, MASS: 4,267) compared with the DL representation (3,796) (Fig. 3c). Additionally, the non-singleton clusters reported by the BIN and MASS representations contained more spectra (BIN: 22,307; 99.68%, MASS: 22,325; 99.76%) than those from the DL representation (16,042; 71.68%), and the adjusted Rand index (ARI) [27] of the clusters reported by the MASS and BIN representations were also higher than that reported by the DL representation (Fig. 3f).

### 2.5 Proteoform identification by spectral library search

We evaluated spectral library search-based proteoform identification using a top-down MS data set described in McCool *et al*. [23], in which 2-dimensional (2D) size exclusion chromatography-capillary zone electrophoresis (SEC-CZE) separation coupled with top-down MS was employed to analyze proteins extracted from SW480 cells. Technical triplicates were included in the data set, and we used the first two replicates, referred to as SW480-2D-1 and SW480-2D-2, in the evaluation. A top-down spectral library was built using the SW480-2D-1 data set with 22,455 MS/MS spectra (see Methods). TopCluster (parameter settings in Supplemental Table S6) was employed to group these spectra into 13,016 spectral clusters. These clusters were further filtered based on proteoform identifications reported by database search using TopPIC [4], reducing the total number of clusters to 5,155. A representative spectrum was computed for each of the 5,155 clusters by averaging the deconvoluted spectra in each cluster (see Methods). The final set of 5,155 representative spectra, corresponding to 3,773 proteoforms, are referred to as the SW480-2D-1 library. To estimate the FDR of identifications, a decoy SW480-2D-1 library of the same size was also built (see Methods).

We searched the 22,924 MS/MS spectra in SW480-2D-2 against the SW480-2D-1 library, combined with the decoy SW480-2D-1 library. Each query spectrum was searched against the combined spectral library to find a matched library representative spectrum with a matched precursor mass and the highest cosine similarity score. Spectral identifications reported from the library search were evaluated using two quality control methods: a 1% spectrum-level FDR and a cosine similarity cutoff of 0.3. Additionally, four combinations of parameter settings were tested, varying the precursor mass error tolerance (10 ppm or 2.2 Da) and the requirement for precursor charge matching (Fig. 4a and 4b). When the cosine similarity cutoff of 0.3 was used as the filtering criterion, the spectrum-level FDR estimated by the target-decoy approach was 0%, and TopLib identified fewer spectra and proteoforms compared with the FDR-based filtering method. As expected, increasing the precursor mass error tolerance and allowing matches between precursors of different charge states resulted in more spectral identifications.

**Fig. 4:**
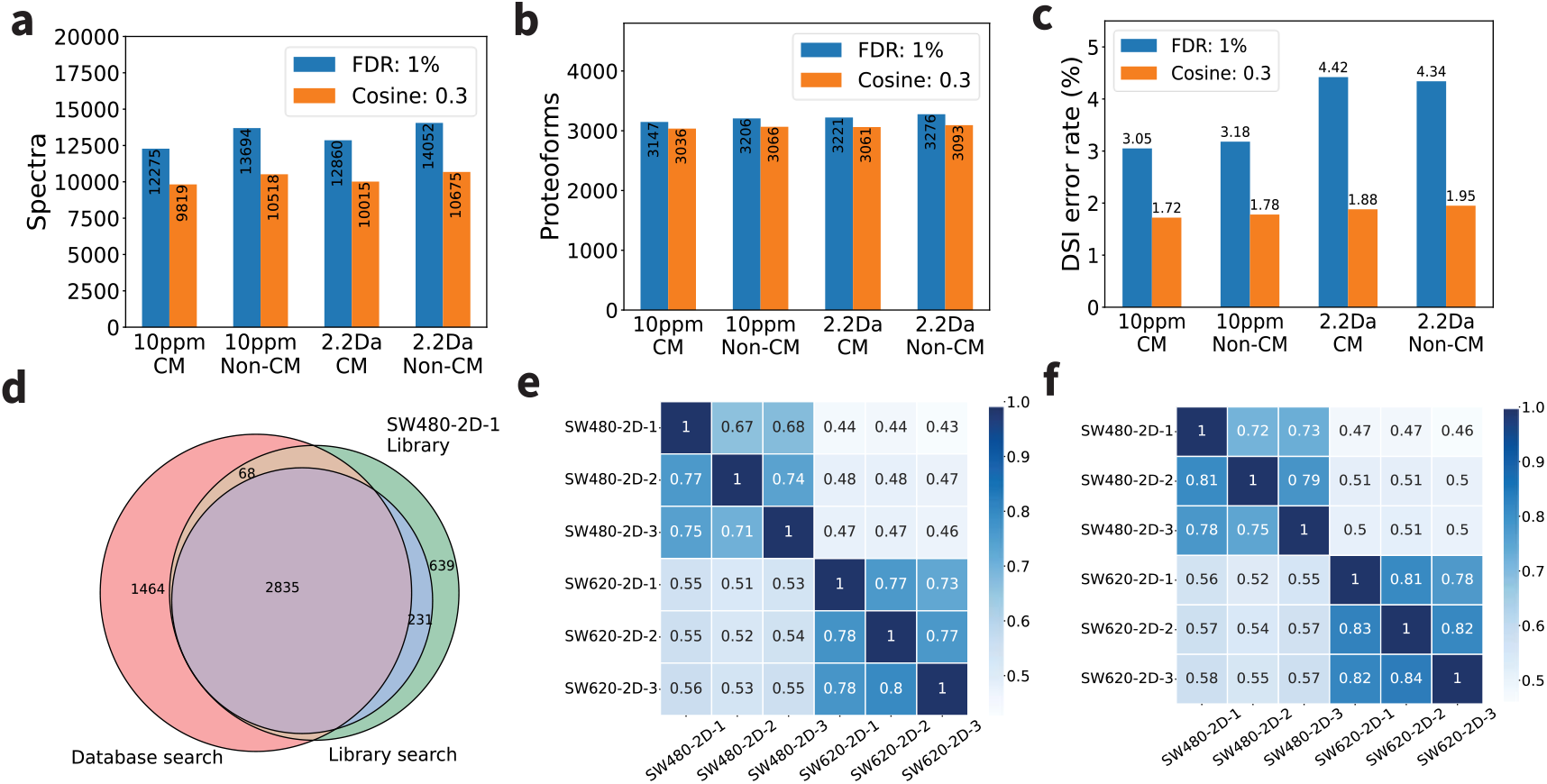
Evaluation of spectral library search. Comparison of (a) spectral identifications, (b) proteoform identifications, and (c) DSI error rates of spectral library search results reported by searching the spectra in SW480-2D-2 against the SW480-2D-1 library with four parameter combinations: a precursor mass error tolerance of 10 ppm or 2.2 Da, and with or without precursor charge matching (CM or Non-CM). Two quality control methods are used: 1% spectrum-level FDR or a cosine similarity cutoff of 0.3 (Cosine: 0.3). (d) Comparison of the proteoforms in the SW480-2D-1 library, the proteoform identifications reported from SW480-2D-2 by database search, and those by spectral library search. Comparison of the reproducibility of proteoform identifications reported by (e) database search and (f) spectral library search.

We evaluated the error rates of spectral library search results based on inconsistent spectral identifications reported by spectral library search and database search. The identifications of a spectrum are inconsistent if the spectrum was matched to two different proteins by the two methods. The error rate of spectral identifications reported by spectral library search was estimated as the ratio of the number of inconsistent identifications to the number of identifications shared by the two search methods, referred to as the database search inconsistency (DSI) error rate. DSI errors may arise from false proteoform annotations in the spectral library, inaccuracies in the SW480-2D-2 database search results, or errors in the spectral library search itself, so the actual error rate for spectral library search is lower than the DSI error rate.

With a 1% spectrum-level FDR, spectral library search produced DSI error rates exceeding 3.05%, suggesting that the FDRs estimated by the target-decoy approach might be underestimated (Fig. 4c). With the cosine similarity-based quality control, the DSI error rates were below 2%. Removing the precursor charge matching requirement increased the number of spectral identifications without significantly affecting the estimated DSI error rates. Increasing the precursor error tolerance resulted in more identifications but also higher estimated error rates. Based on these findings, we chose the default parameter settings for spectral library search in TopLib as follows: 10 ppm for the precursor error tolerance, no requirement for precursor charge matching, and a cosine similarity cutoff of 0.3.

### 2.6 Comparison between database search and spectral library search

We compared spectrum and proteoform identifications reported from SW480-2D-2 by database search using TopPIC (version 1.7.5, parameter settings in Supplemental Table S5) and by spectral library search using TopLib with the default parameter settings. TopLib identified 231 proteoforms and 1,128 spectra missed by database search (Fig. 4d and Supplemental Fig. S6). This improvement is attributed to the inclusion of mass signal intensity information in the library spectra, which enhances the sensitivity of spectral library search in comparison with database search. On the other hand, TopLib missed 2,738 spectra and 1,532 proteoforms that were identified by database search. The primary reason for these missed identifications was the incompleteness of the spectral library: 1,464 (95.6%) of the 1,532 missed proteoforms were due to missing library spectra. Of the remaining 68 missed identifications, 48 were due to large errors in precursor masses, and 20 were due to low MS/MS spectral similarity.

TopLib was also 140 times faster than database search for spectral identification. On a computer with an 11th Gen Intel Core <u>i9-11900K 3.5GHz</u> CPU and 64 GB of memory, the running time of the spectral library search with one CPU thread was 3.35 minutes, whereas the running time of the database search with 16 CPU threads was 470 minutes.

We also evaluated the reproducibility of proteoform identifications reported by database search using TopPIC and by spectral library search using TopLib across six SEC-CZE data sets (three from SW480 cells and three from SW620 cells) described by McCool et al. [23]. The first two data sets, referred to as SW480-2D-1 and SW480-2D-2, were used in the previous section, while the other four are referred to as SW480-2D-3, SW620-2D-1, SW620-2D-2, and SW620-2D-3. We searched the MS/MS spectra in the six data sets against the UniProt human proteome database to identify spectra and proteoforms using TopPIC (version 1.7.5, parameter settings in Supplemental Table S5). For each data set pair *A* and *B* in the six data sets, the reproducibility of proteoform identifications for database search was computed as the ratio between the number of identifications shared by the two data sets and the number of identifications reported from *A*. The reproducibility of proteoform identifications for spectral library search was obtained using the following method. We built a spectral library using data set *A* (see Methods) and searched the MS/MS spectra in data set *B* against the spectral library to identify spectra and proteoforms using TopLib with the default parameter settings. The reproducibility of proteoform identifications for the spectral library search was the ratio between the number of proteoform identifications reported by the spectral library search and the number of proteoform identifications reported from *A* by the database search. On average, spectral library search improved the reproducibility of proteoform identifications by 2.63% compared with database search (Fig. 4e and 5f).

## 3. Methods

### 3.1 SQL database design

An SQLite database is used to store MS/MS spectra and proteoform identifications of top-down MS data in TopLib. The relational diagram depicting the tables of the database is given in Supplemental Fig. S7. The precursor information and deconvoluted fragment masses of MS/MS spectra are stored in a spectrum and a mass table, respectively. Representative spectrum tables are used to store the information of representative spectra.

### 3.2 MS data preprocessing and database search

The raw MS data files were converted to mzML files using msconvert (version 3.0.10765) [28] and deconvoluted to msalign files using TopFD (version 1.7.5 and see Supplemental Table S4 for parameter settings) [14]. These msalign files were subsequently searched against the UniProt human proteome sequence database (UP000005640_9606, 20,590 entries, version July 19, 2024) concatenated with a decoy database of the same size using TopPIC (version 1.7.5 and see Supplemental Table S5 for parameter settings) [4].

### 3.3 Evaluation data sets

In data preprocessing, TopFD grouped precursor ions in each data file into proteoform features. The precursor ions in each proteoform feature had similar molecular masses and similar elution times in proteoform separation. All PrSMs reported from the SW480-3D data set by database search were divided into proteoform groups using proteoform features reported by TopFD and proteoform identifications reported by TopPIC: Two PrSMs were assigned to the same proteoform group if the precursor ions of the two MS/MS spectra were from the same proteoform feature reported by TopFD or the two PrSMs were matched to the same protein and their precursor mass difference was no more than 10 ppm.

Because spectral deconvolution may introduce ±1, ±2 Da errors into precursor masses of proteoforms, we removed possible duplicated proteoform groups as follows. All the proteoform groups were first ranked in the increasing order of the *E*-value based on their best PrSMs. For a proteoform group *A* matched to a protein *P*, if we found another proteoform group *B* such that (1) *B* was ranked higher than *A*, (2) *B* matched to protein *P*, and (3) the precursor mass difference between *A* and *B* was less than 2.2 Da, then the proteoform group *A* was removed.

We further removed PrSMs with inconsistent identifications. Two PrSMs were inconsistent if the two spectra were assigned to the same proteoform group, but they were matched to two different proteoforms. Note that these inconsistent PrSMs were assigned to the same proteoform group because their precursor ions were from the same proteoform feature reported by TopFD even though their proteoform identifications reported by database search were different. To remove inconsistent identifications, we ranked all PrSMs in the same proteoform group in the increasing order of the *E*-value. For a PrSM *A*, if we could find another PrSM *B* in the same proteoform group such that (1) *B* had a better *E*-value than *A* and (2) PrSMs *A* and *B* were matched to two different proteins, then PrSM *A* was removed.

To build the SW480-SPE data set, we shifted fragment masses in the spectra of the same-group spectral pairs. For each same-group spectral pair (*S*_1_, *S*_2_), spectrum *S*_1_ remained unchanged, and 90% of the fragment masses in *S*_2_ were shifted by random values within the ranges of [-200, -100] or [100, 200] Da. If a shifted mass was below zero or exceeded the precursor mass, a new random value within the ranges was selected to ensure that the shifted mass remained between 0 and the precursor mass.

To generate the SW480-GPSE data set, for a given charge state *c*, we selected all SW480-3D spectral groups with the charge state *c*. For each spectral group, the spectrum with the best *E*-value PrSM was chosen as the representative spectrum. Then we ranked the representative spectra using their precursor masses in the decreasing order. For *i* = 1, 3, 5, …, we calculated the precursor mass difference between the *i*th and *i*+1th representative spectra, and then increased the precursor masses of all spectra in the *i*+1th group by the mass difference.

### 3.4 Mass spectral representations

Spectral similarity scores and distances were calculated using deconvoluted top-down mass spectra containing neutral monoisotopic masses and intensities. Three representation methods were employed to convert a deconvoluted mass spectrum into a vector of real numbers to speed up spectral similarity/distance computation. For each spectrum, only the *k* most intense (mass, intensity) pairs were retained to simplify the data and the default setting for *k* was set to 50. In the DL representation, each mass in a spectrum is converted to its corresponding monoisotopic *m/z* value, the intensities are normalized to a unit length, and a deep neural network [12] is utilized to encode the (*m*/*z*, intensity) pairs to a vector of size 32. In the BIN and MASS representations, log transformation (base 2) is applied to the intensities, and the log-transformed intensities are normalized to a unit length. In the BIN representation, masses are also converted to their corresponding monoisotopic *m/z* values, and a bin-based method, with a user-specified bin size, is utilized to convert the (*m*/*z*, log(intensity)) pairs in the *m*/*z* range of [200, 1700] into a vector of log intensities [11], which is further normalized to a unit length. In the MASS representation, the spectrum is represented by its mass and normalized intensity pairs.

### 3.5 Distance functions for mass lists

In the MASS representation, two spectra *S* and *T* are represented as lists of (mass, intensity) pairs, in which the mass intensities in each spectrum are normalized to a unit length. An intensity *x* from *S* is matched to an intensity *y* is from *T* if the distance between their corresponding mass values is less than an error tolerance. Additionally, ±1.00235 Da errors are allowed in matching fragment masses. If an intensity in *S* or *T* cannot be matched to any intensity in the other spectrum, it will be paired with a zero intensity. Euclidean distance, cosine distance, and the entropy-based distance of the mass representations of *S* and *T* are defined on the two normalized unit vectors obtained from the paired intensities. To speed up the distance computation in TopLib, the mass intensities in *S* and *T* are normalized to a unit length before the paired intensities are found, and spectral distances are computed using these normalized intensities. Because one mass in one spectrum may be matched to multiple masses in the other, the normalized intensities computed based on paired intensities may be slightly different from those calculated based on single spectra.

### 3.6 Top-down spectral clustering

In TopCluster, mass spectra are clustered in three steps. (1) All spectra are grouped based on their precursor charge state, ensuring that the spectra in a group share the same charge state. (2) The spectra in each group reported from step (1) are clustered based on their precursor masses: each spectrum is represented by only its precursor masses and clustered using hierarchical clustering with the complete linkage and a distance threshold of 2.2 Da. (3) In the final step, the spectral clustered reported from step (2) are further clustered using the fragment masses in their MS/MS spectra. Pairwise spectral cosine distances are calculated and then used as input for hierarchical clustering with average linkage or DBSCAN clustering [29].

### 3.7 Evaluation metrics for spectral clustering

Clustering performance were evaluated using ARI [27], clustered spectra ratio, incorrect clustering ratio, and completeness [11, 12, 30]. For two partitions *A* and *B* of the same set of spectra, ARI is computed as follows:

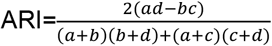

where *a* is the number of spectral pairs placed in the same cluster by both partitions *A* and *B*; *b* is the number of pairs placed in the same cluster in partition *A*, but in different clusters in partition *B*; *c* is the number of pairs placed in the same cluster in partition *B* but in different clusters in partition *A*; and *d* is the number of pairs of placed in the different clusters by both partitions *A* and *B*.

A cluster with at least two spectra is called a valid cluster, and any spectrum assigned to a valid cluster is considered as a clustered spectrum. *The ratio of clustered spectra* of a partition of a set of spectra is the fraction of clustered spectra in the entire set.

Let *A* be the ground truth partition of a spectral set and *C* is a valid cluster reported by a clustering method from the spectral set. We further divide *C* into sub-clusters based on the clusters in *A*: two spectra in *C* are assigned to the same sub-cluster if they are in the same cluster in *A*. The spectra in the largest sub-cluster are considered as correctly clustered ones; the remaining ones are incorrectly clustered. If multiple sub-clusters contain the same largest number of spectra, one is randomly selected as the “largest” one. *The ratio of incorrectly clustered spectra* of a partition is the ratio between the numbers of incorrectly clustered spectra and clustered spectra.

Consider a set of *N* spectra with a ground truth partition *A*={*A*_1_, *A*_2_, …, *A*_*K*_} with *K* clusters and another partition *B =*{*B*_1_, *B*_2_, *…, B*_*L*_} with *L* clusters of. Let *n*_*j*_ be the total number of spectra in *B*_*j*_ and *n*_*i,j*_ the number of spectra shared by *B*_*i*_ and *A*_*j*_. The completeness of *B* with respect to *A* is defined as 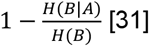, where

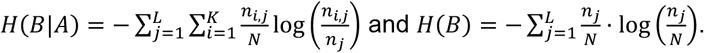

### 3.8 Building spectral libraries

Raw MS files were preprocessed using msconvert [28] TopFD [14] and deconvoluted spectra were identified by database search using TopPIC [4] (see Section 3.2). MS/MS spectra lacking deconvoluted precursor information or containing fewer than two fragment masses were discarded. The remaining spectra were clustered using TopCluster (parameter settings in Supplemental Table S6).

Possible incorrect proteoform identifications reported by TopPIC were filtered following the method described in Section 3.3. Spectral clusters without any proteoform identifications reported by TopPIC were discarded. If the spectra in a cluster are matched several proteoforms, the proteoform with the best E-value PrSM reported by TopPIC was selected for the cluster. If two clusters were matched to the same proteoform with the same charge state, only the cluster with the best E-value PrSM was retained, and the other was removed.

### 3.9 Representative spectra

Two types of representative spectra were generated for a spectral cluster: single representative spectra and average representative spectra. Given a spectral cluster, if some spectra in the cluster had proteoform identifications reported by database search, the spectrum corresponding to the best E-value PrSM was selected as the single representative spectrum. Otherwise, the spectrum with the largest number of fragment masses was selected as the single representative spectrum.

To generate the average representative spectrum of a cluster, we first computed the merged spectrum *T* of two spectra in the cluster, and then the merged spectrum of *T* and the third spectrum. This process was repeated until the merged spectrum of all the spectra in the cluster was obtained. An error tolerance of 10 ppm was used to merge deconvoluted fragment masses in the spectra. For two masses *x*_1_ and *x*_2_ with their corresponding intensities *y*_1_ and *y*_2_, the merged mass was their weighted average mass *x* = (*x*_1_*y*_1_ + *x*_2_*y*_2_)/(*y*_1_+*y*_2_). Finally, the 50 most intense fragment masses in the merged spectrum were selected, their intensities were normalized to a unit length, and the resulting (mass, intensity) pairs were reported as the representative spectrum.

### 3.10 Decoy spectral libraries

For each representative spectrum in the spectral library, a decoy spectrum is generated using a mass shifting method. For each fragment mass in the representative spectrum, two different amino acids are randomly selected, and the fragment mass is shifted by the difference between their monoisotopic masses. If the shifted mass falls below zero or exceeds the precursor mass, a new random shift is considered until the shifted mass is between 0 to the precursor mass.

## 4. Conclusions and Discussion

We developed TopLib, the first software tool for building and searching top-down spectral libraries for proteoform identification. TopLib leverages fragment mass signal intensities in top-down MS/MS spectra to enhance the sensitivity of spectral identification. As a result, TopLib improves the reproducibility of proteoform identifications compared with database search-based methods. Additionally, TopLib is 140 times faster than TopPIC [4], a commonly used database search tool for top-down mass spectral identification.

We compared the performance of three spectral representation methods: DP, BIN, and MASS, and found that the MASS representation outperformed the other two in distinguishing between correct and incorrect SSMs. Additionally, applying log transformation to mass signal intensities enhanced the ability to distinguish between correct and incorrect SSMs. By using deconvoluted centroided fragment masses, we simplified the representation of top-down mass spectra and demonstrated that spectra containing the 50 most intense fragment masses achieved similar performance in identifying correct SSMs compared with spectra with all deconvoluted fragment masses. The MASS representation also resulted in higher accuracy in spectral clustering than the other representation methods. Furthermore, for the MASS representation, no significant differences were observed across Euclidean distance, cosine distance, and the entropy-based distance.

We also comprehensively evaluated the performance of TopLib for spectral identification with various parameter settings. Using inconsistent identifications reported by database search and TopLib, we found that the FDR estimation based on target and decoy spectra may underestimate the error rate of the identifications reported by spectral library search. To address this issue, further research is needed to explore alternative methods for generating decoy spectra and estimating FDRs. In addition, removing the requirement for precursor charge state matching can increase spectral identifications without significantly increasing the error rate.

TopLib still has several limitations. First, it relies on comprehensive spectral libraries for spectral identification, restricting its applications in discovery-mode studies. Second, TopLib currently does not support querying spectra with unexpected mass shifts. Third, library spectra in TopLib are not fully annotated. Since TopLib uses PrSMs reported by database search to build spectral libraries, some library spectra may correspond to proteoforms with unknown mass shifts. Manual inspection and new proteoform characterization methods are needed to enhance spectral annotation in these libraries.

## Code availability

The source code of TopLib is available at https://github.com/toppic-suite/toplib.

## Acknowledgements

This research was funded by NIH through the grants R01GM118470 and R01CA247863.

## Language polishing

The authors used ChatGPT to enhance the language and readability during the preparation of this paper. After utilizing ChatGPT, the authors reviewed and edited the content and take full responsibility for the final version of the paper.

