## Supplemental Material for "TopLib: Building and searching top-down mass spectral libraries for proteoform identification"

1. **Supplemental Figures**

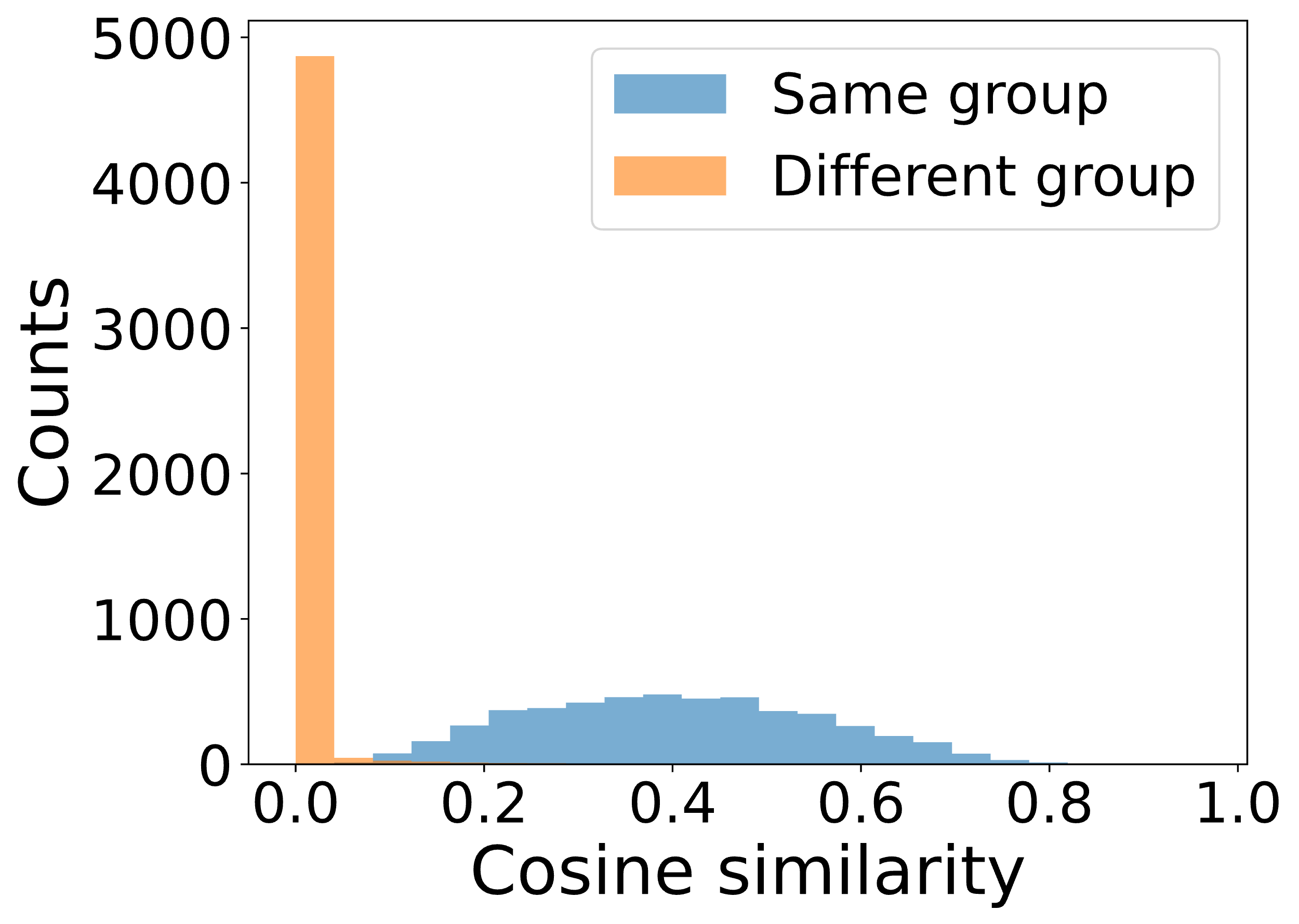

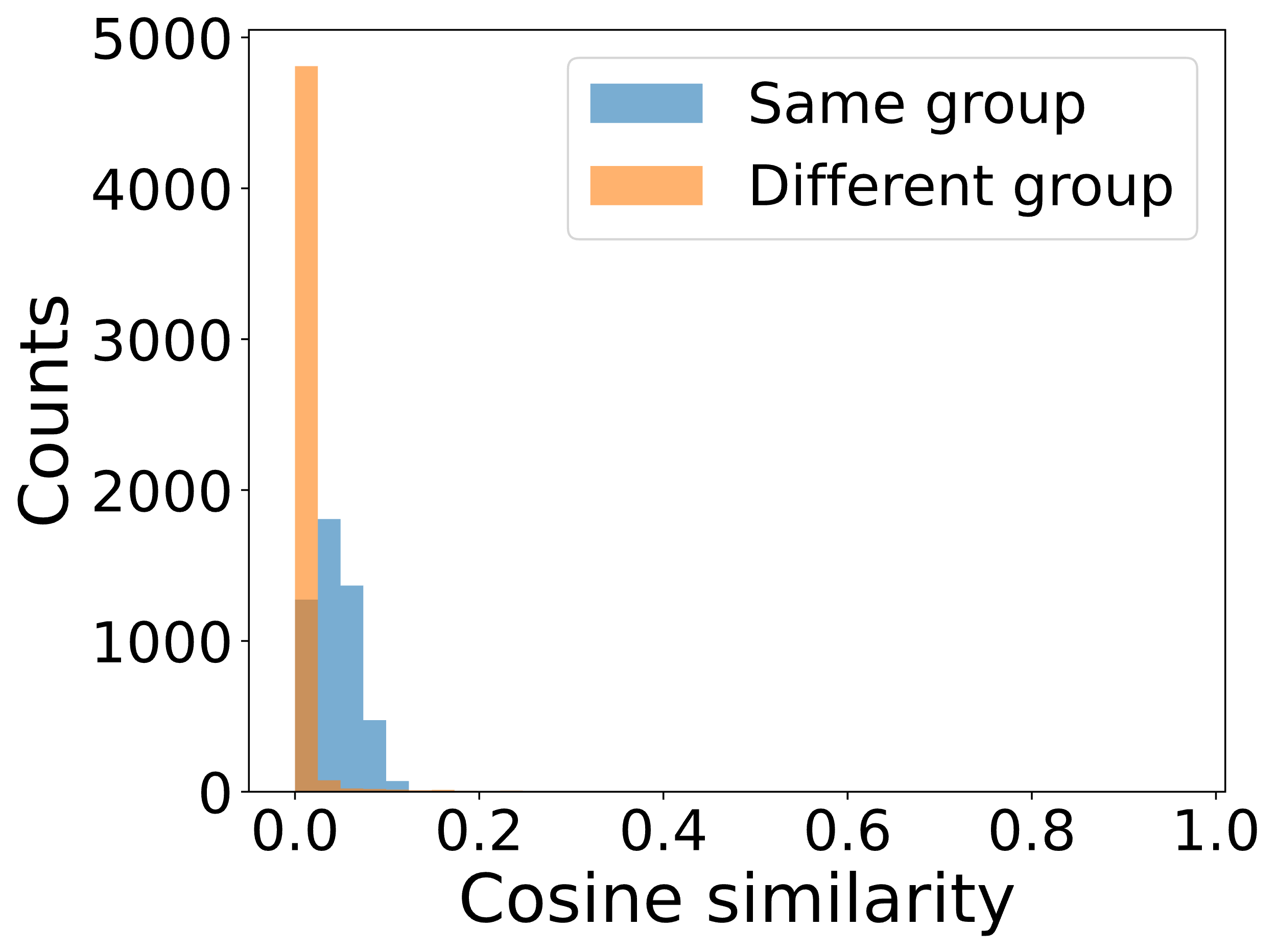

(a)	(b)

**Fig. S1:** The distributions of cosine similarity of same-group and different-group spectrum pairs in the SW480-SPE data set without mass shifts (a) and with shifts for 90% of the masses in spectra (b).

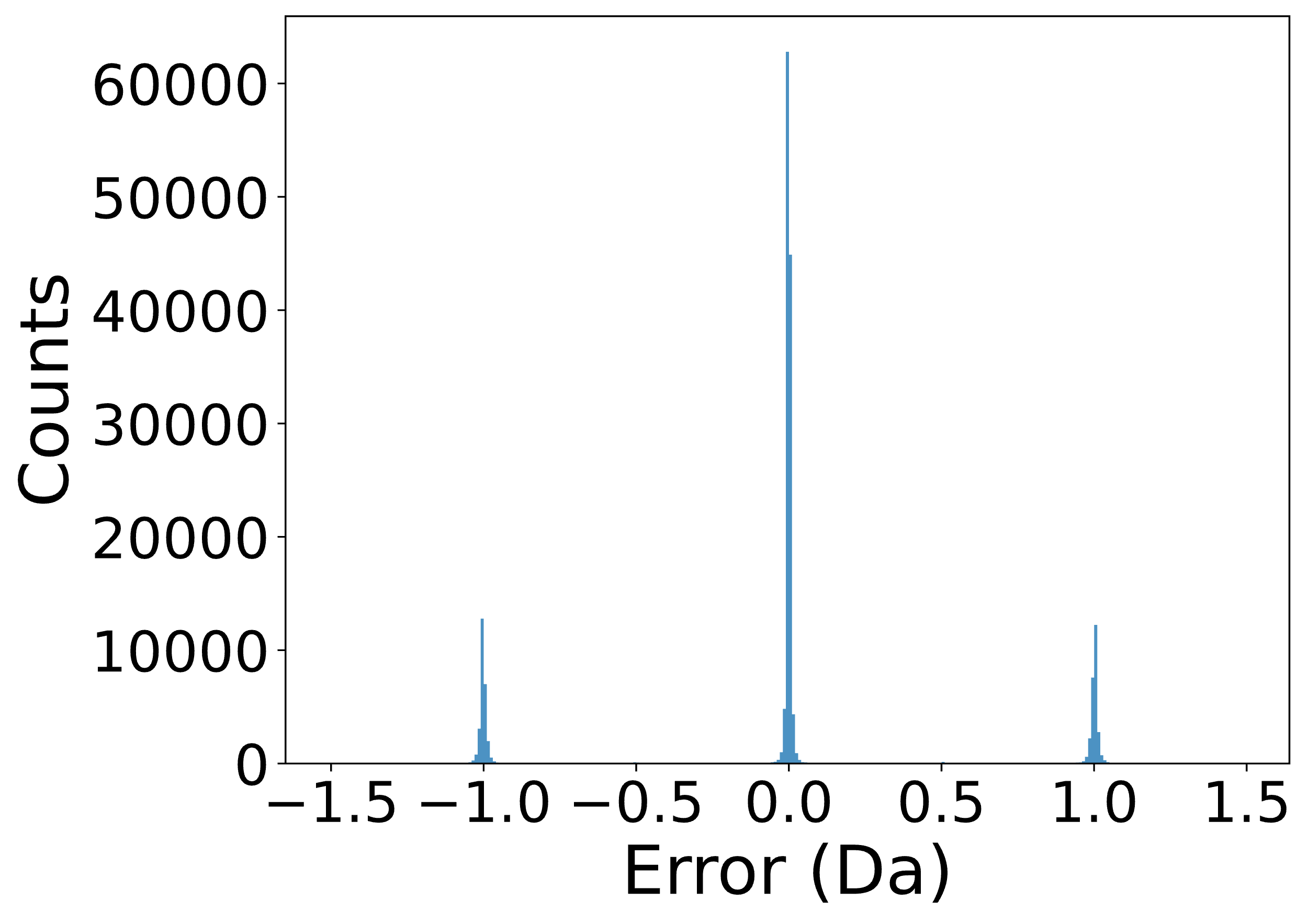

**Fig. S2:** The distribution of the errors of matched fragment masses in the same-group spectrum pairs in the SW480-SPE data set.

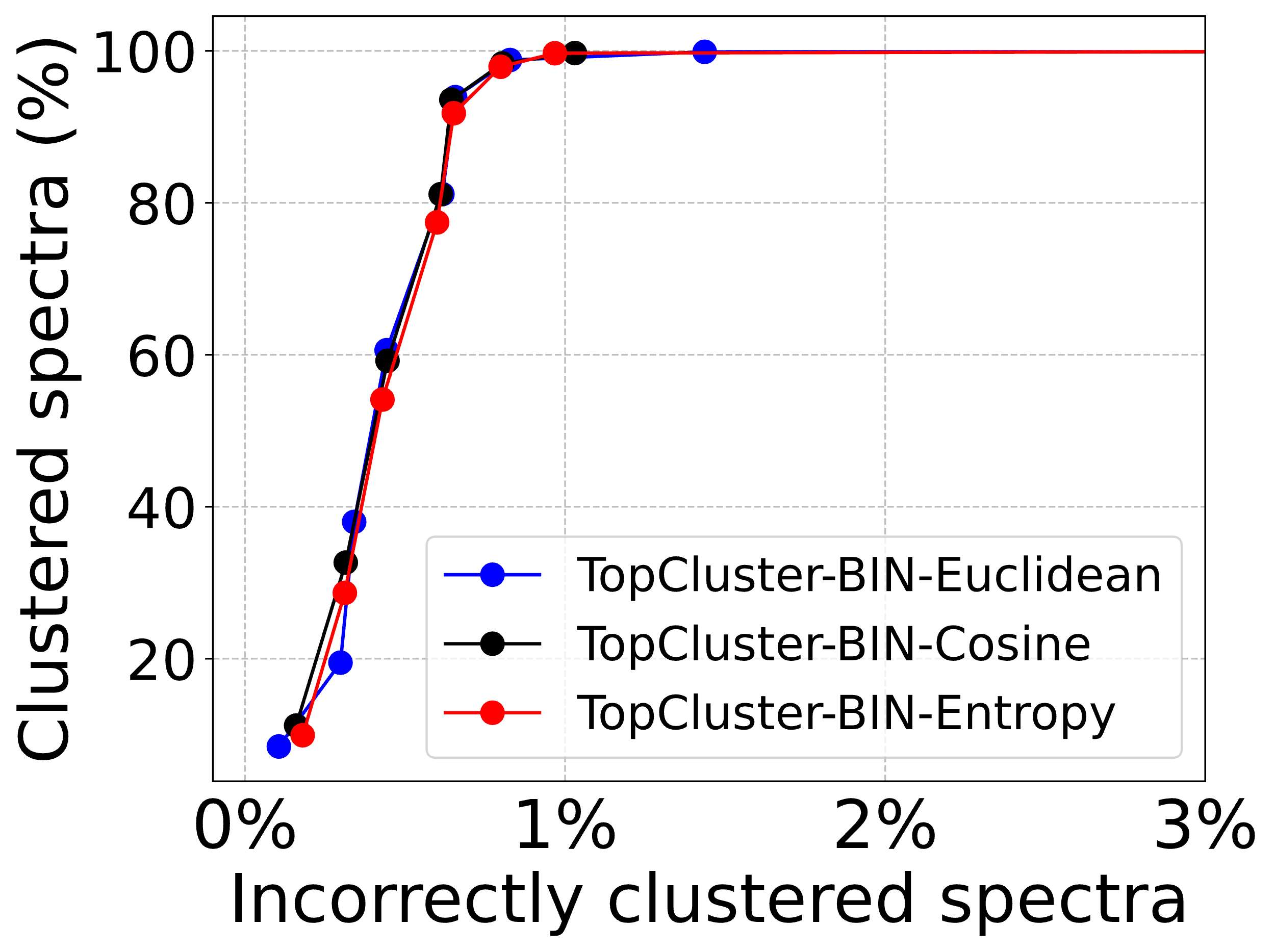

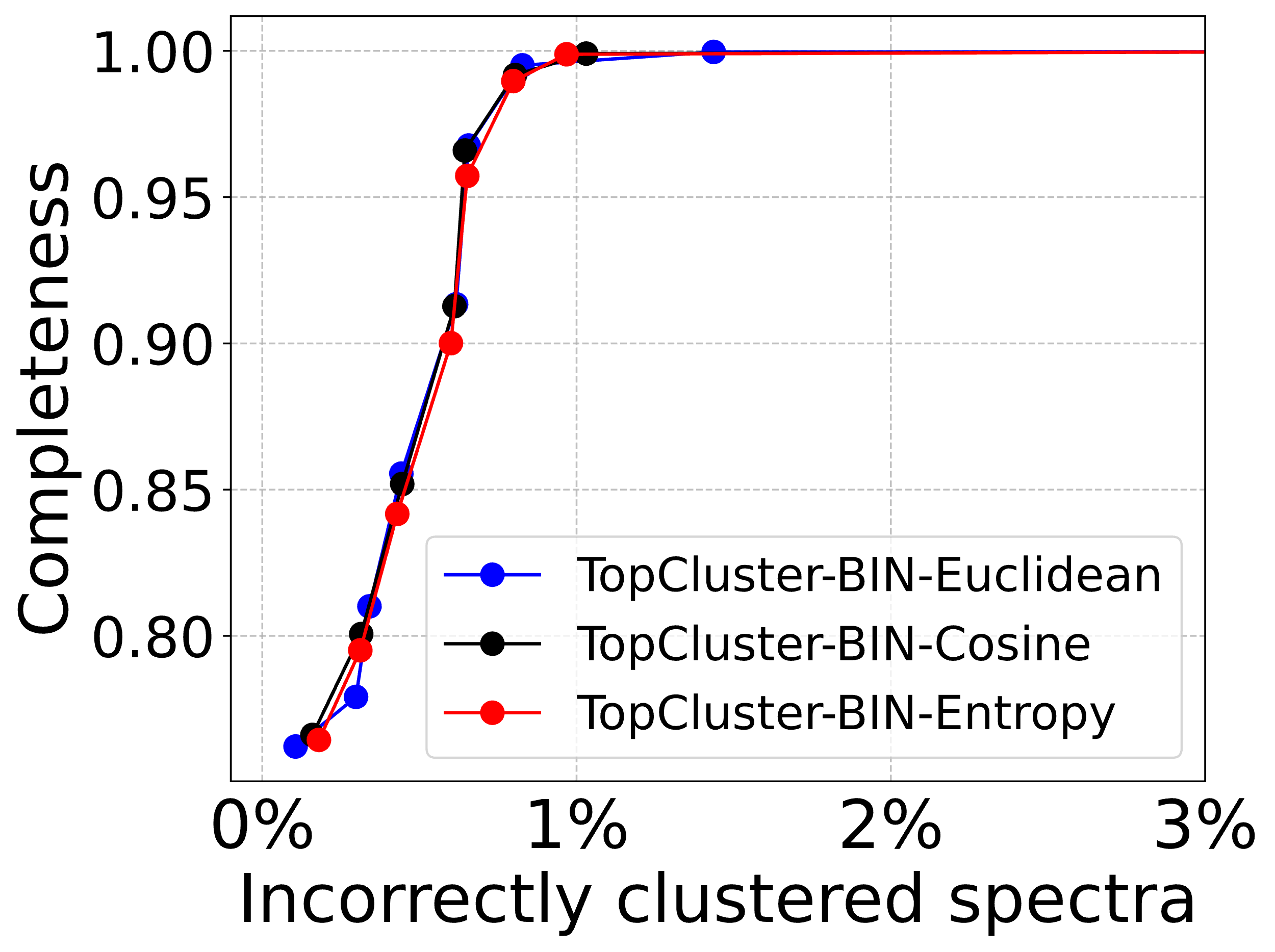

(a)	(b)

**Fig. S3:** Comparison of spectral clustering accuracy of TopCluster with the BIN representation and hierarchical clustering using three distance functions: Euclidean, Cosine, and the entropy-based distances. (a) The ratio of incorrectly clustered spectra against the ratio of clustered spectra. (b) The ratio of incorrectly clustered spectra against clustering completeness.

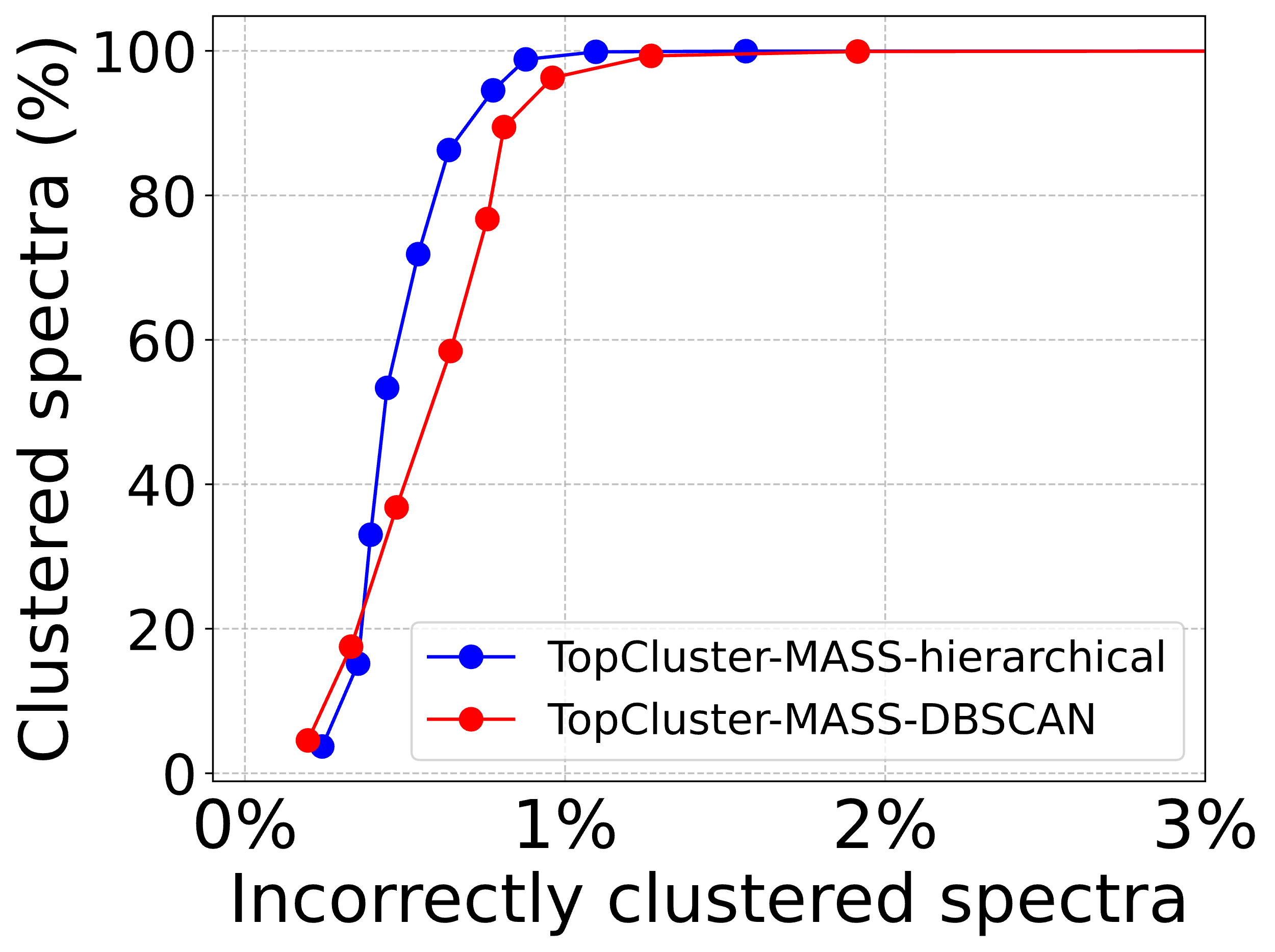

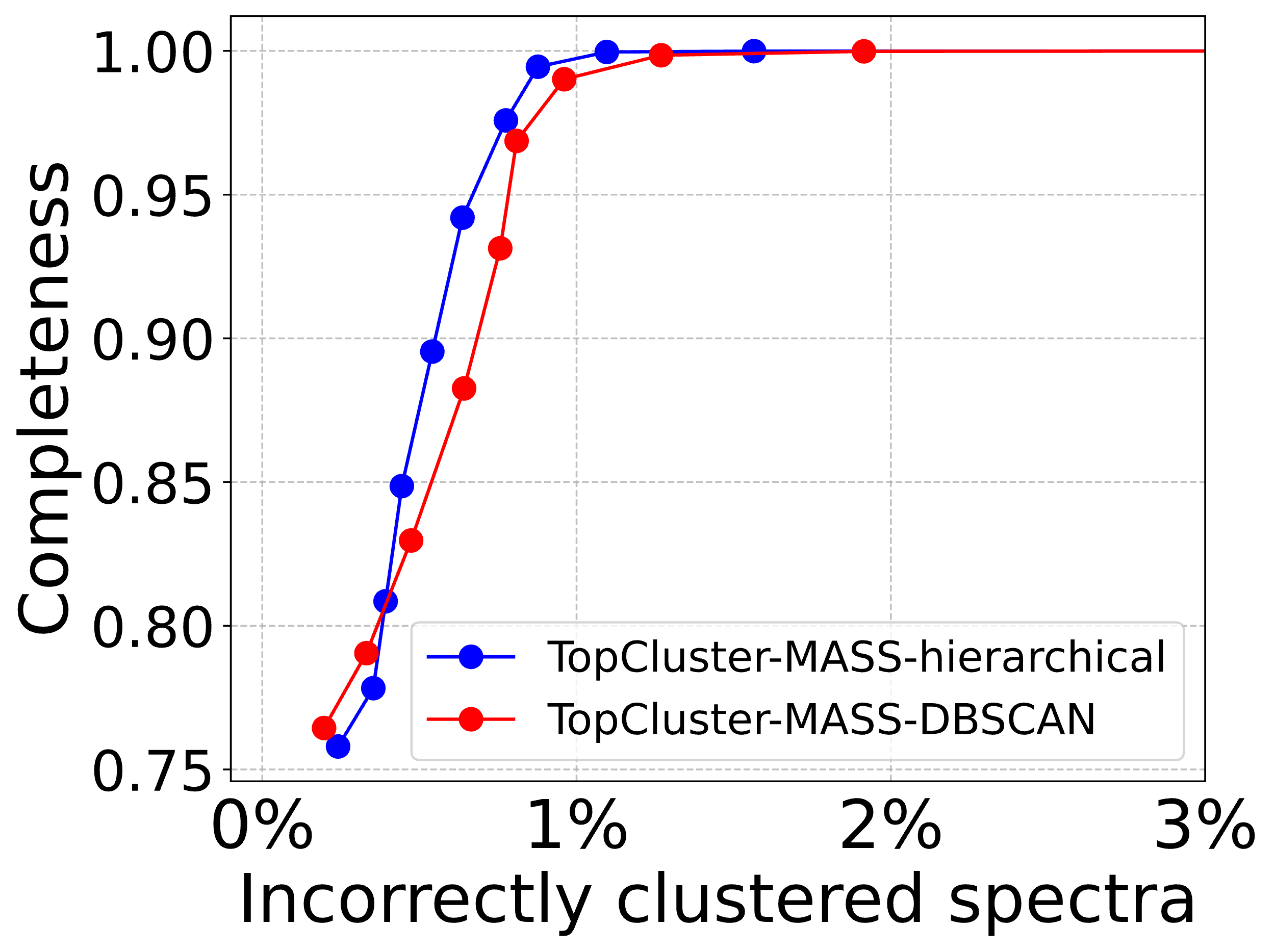

(a)	(b)

**Fig. S4:** Comparison of spectral clustering accuracy between hierarchical clustering and DBSCAN using TopCluster with the MASS representation. (a) The ratio of incorrectly clustered spectra against the ratio of clustered spectra and (b) the ratio of incorrectly clustered spectra against clustering completeness.

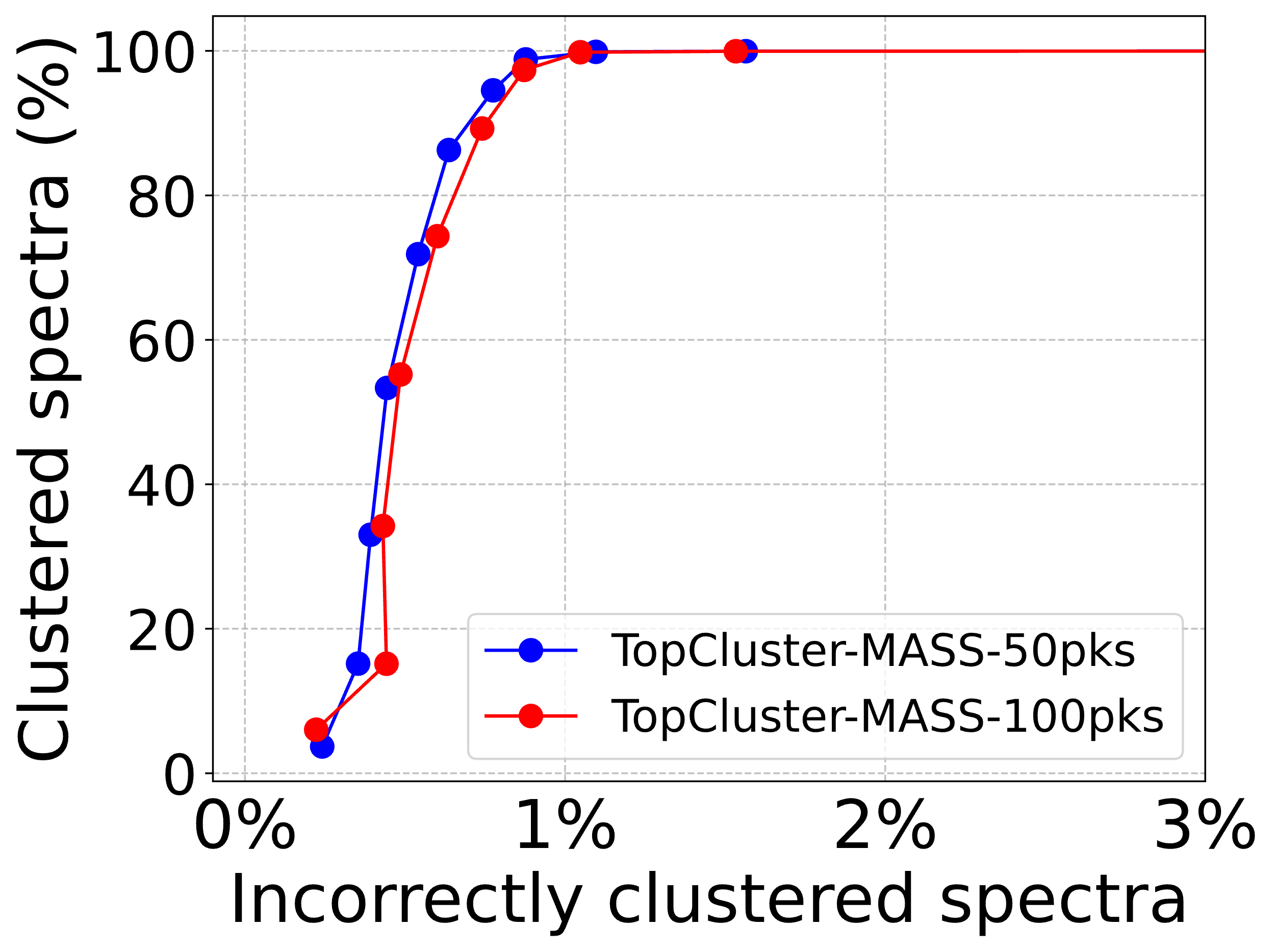

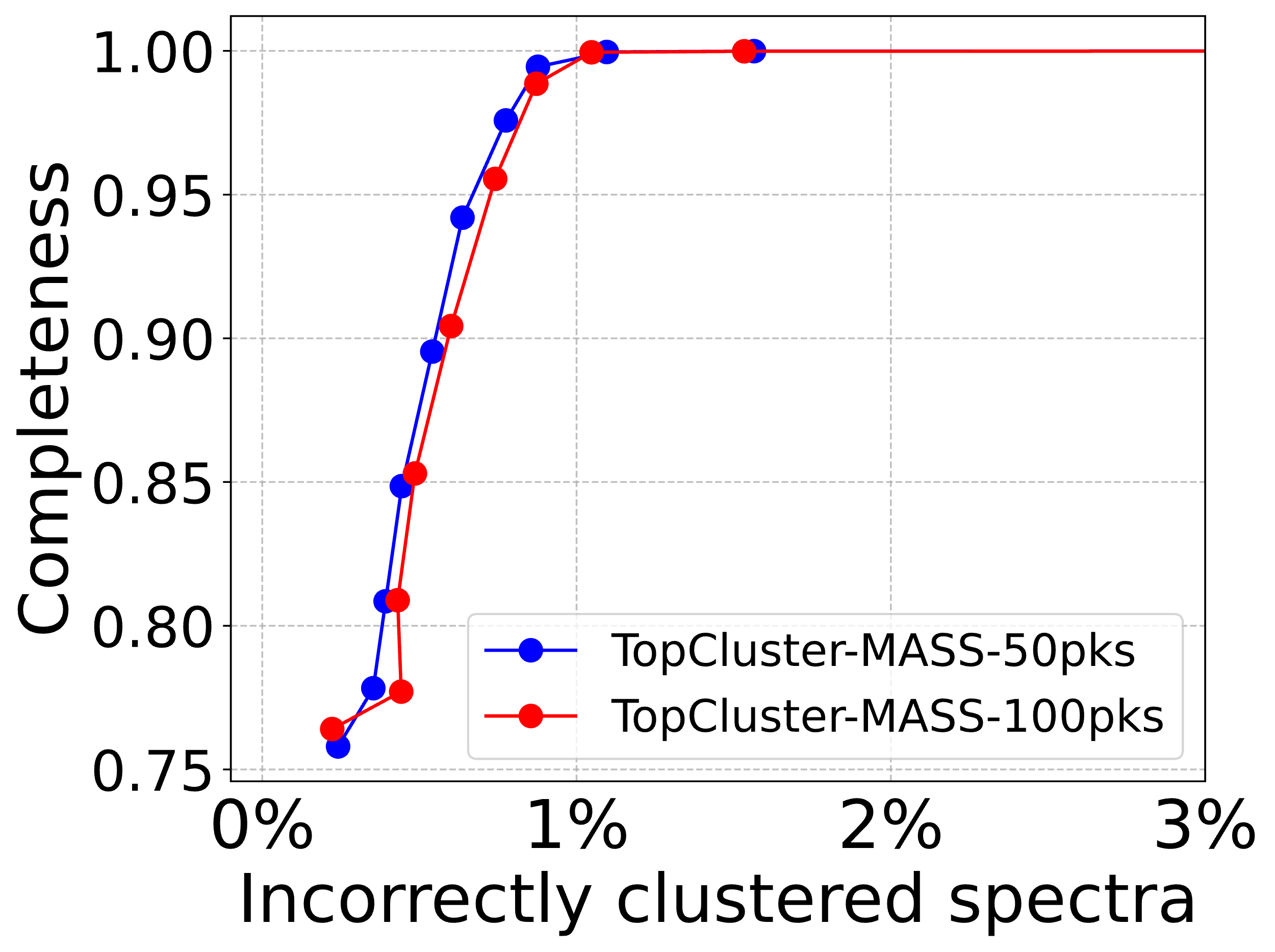

(a)	(b)

**Fig. S5:** Comparison of spectral clustering performance between *k*=50 and *k*=100 for TopCluster using the MASS representation. (a) The ratio of incorrectly clustered spectra against the ratio of clustered spectra and (b) the ratio of incorrectly clustered spectra against clustering completeness.

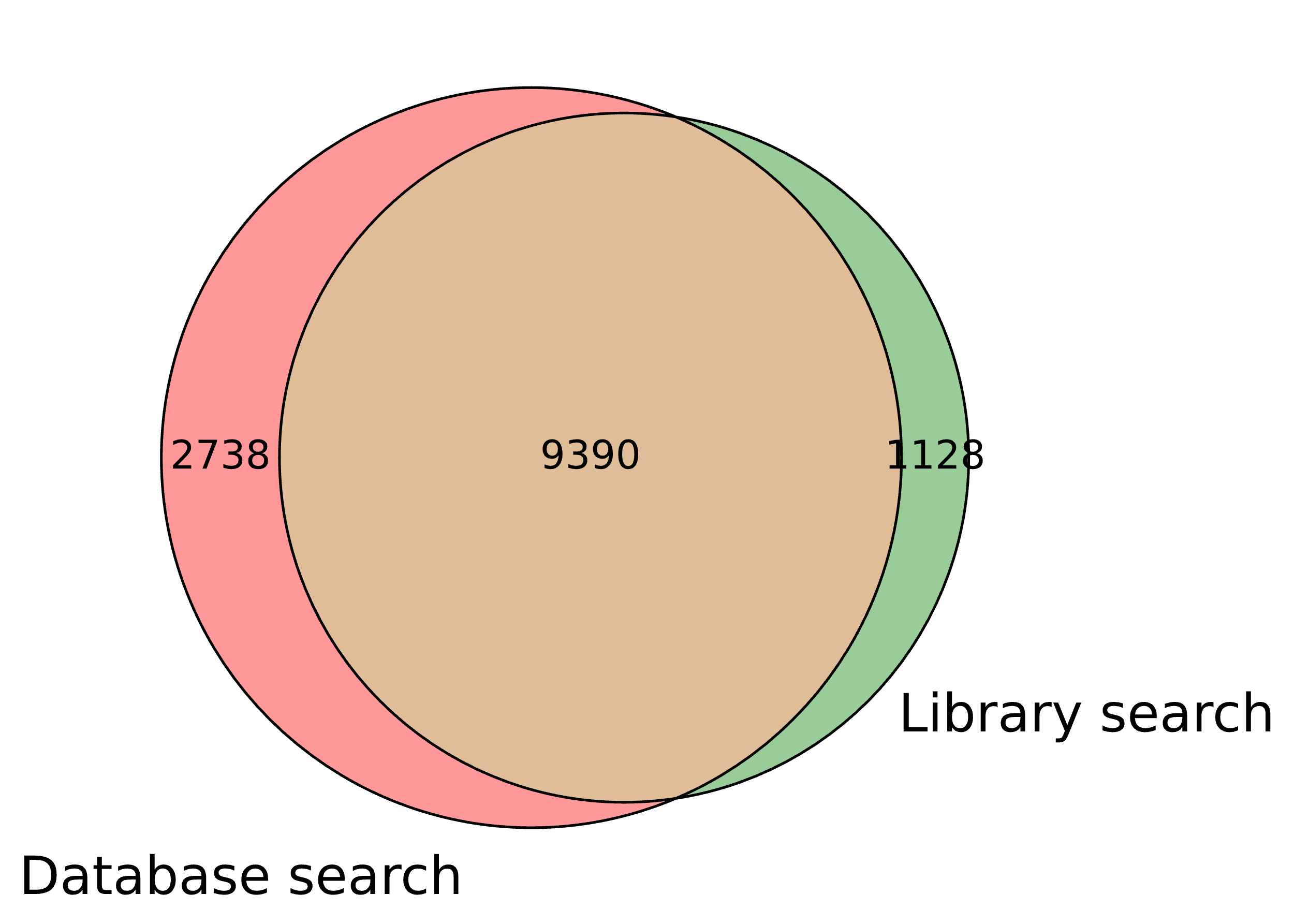

**Fig. S6:** Comparison of spectral identifications reported from SW480-2D-2 by database search and spectral library search against the SW480-2D-1 library.

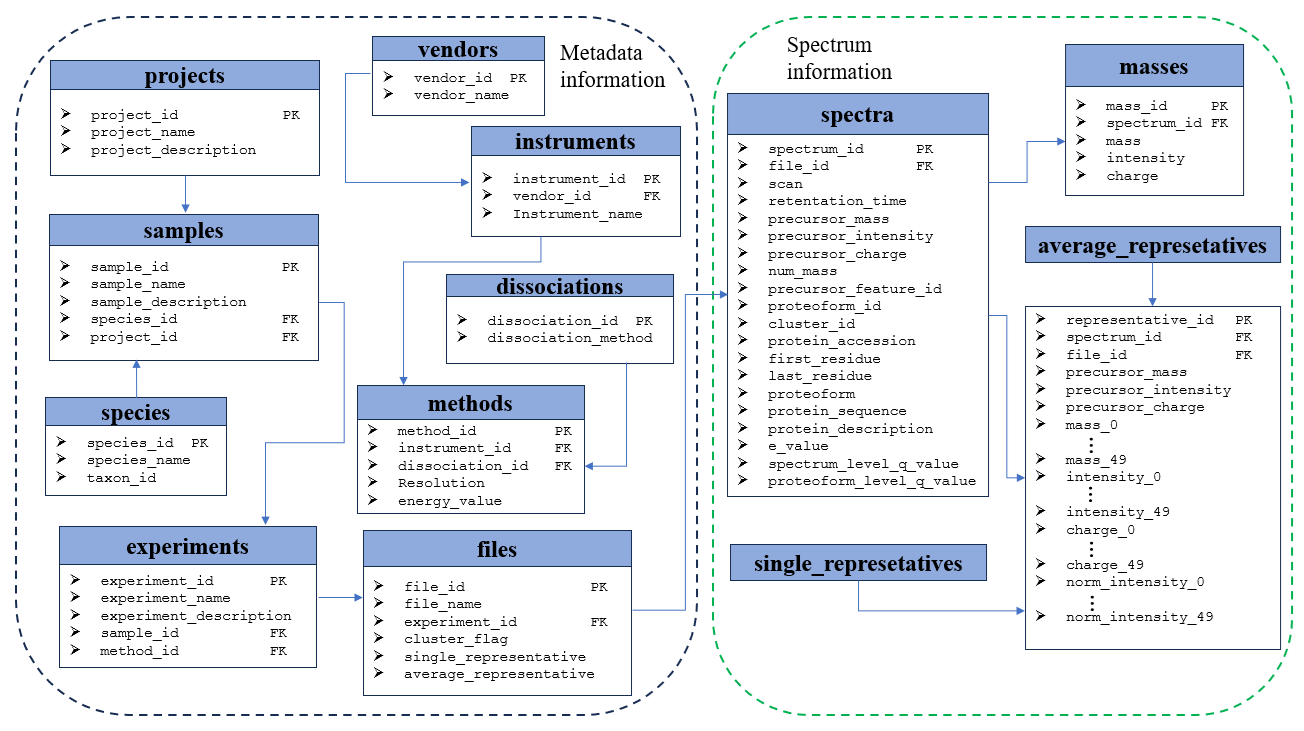

**Fig. S7:** Database scheme diagram for storing MS data in TopLib.

2. **Supplemental Tables**

**Table S1:** Comparison of different settings of *k* using the BIN representation on the SW480-SPE data set with a bin size of 0.5

| Bin=0.5 | *k*=25 | *k*=50 | *k*=75 | *k*=100 | *k*=150 | *k*=200 |
| --- | --- | --- | --- | --- | --- | --- |
| Euclidean | 0.85414 | **0.87251** | 0.85336 | 0.83435 | 0.80603 | 0.79586 |
| Cosine | 0.85500 | **0.87244** | 0.85335 | 0.83435 | 0.80603 | 0.79585 |
| Entropy | 0.85125 | **0.87107** | 0.86367 | 0.85390 | 0.83858 | 0.83327 |

**Table S2:** Comparison of different settings of *k* using the DL representation on the SW480-SPE data set

|  | *k*=25 | *k*=50 | *k*=75 | *k*=100 | *k*=150 | *k*=200 |
| --- | --- | --- | --- | --- | --- | --- |
| Euclidean | 0.59197 | **0.59860** | 0.59215 | 0.58266 | 0.57427 | 0.57218 |
| Cosine | 0.59795 | **0.61585** | 0.61125 | 0.60834 | 0.60589 | 0.60646 |

**Table S3:** Comparison of different settings of *k* using the MASS representation on the SW480-SPE data set with an error tolerance of 10 ppm

|  | *k*=25 | *k*=50 | *k*=75 | *k*=100 | *k*=150 | *k*=200 |
| --- | --- | --- | --- | --- | --- | --- |
| Euclidean | 0.87385 | 0.92455 | 0.92871 | **0.93154** | 0.93059 | 0.92920 |
| Cosine | 0.87408 | 0.92306 | 0.93020 | **0.93178** | 0.93016 | 0.92956 |
| Entropy | 0.87529 | 0.92571 | 0.93391 | **0.93618** | 0.93533 | 0.93507 |

**Table S4:** Parameter settings for TopFD

| **Parameter** | **Value** |
| --- | --- |
| Spectral data type | Centroid |
| Maximum charge | 30 |
| Maximum monoisotopic mass | 70000 Dalton |
| Peak error tolerance | 0.02 m/z |
| M1 signal/noise ratio | 3 |
| MS/MS signal noise ratio | 1 |
| Precursor window | 3 m/z |
| Do final filtering | Yes |

**Table S5.** Parameter settings for TopPIC

| Parameter | Value |
| --- | --- |
| Proteome database | UniProt human proteome database (UP000005640_9606, 20,590 entries, version July 19, 2024) |
| Fragmentation method | File |
| Search type | Target+Decoy |
| N-terminal forms of proteins | NONE, M_ACETYLATION, NME, NME_ACETYLATION |
| Use TopFD features | Yes |
| Fixed modifications | No |
| Maximum number of variable modifications | 0 |
| Variable modifications | None |
| Spectrum level cutoff type for filtering PrSMs | FDR |
| The cutoff value for filtering PrSMs | 0.01 |
| Spectrum level cutoff type for filtering proteoforms | FDR |
| The cutoff value for filtering proteoforms | 0.01 |
| Error tolerance for precursor and fragment masses | 10 ppm |
| Error tolerance for identifying PrSM clusters | 10 ppm for building spectral library;  2.2 Da for estimating error rates of spectral library search. |
| Maximum number of unexpected mass shifts | 1 |
| Minimum value of the mass shift | -500 |
| Maximum value of the mass shift | 500 |
| E-values computation | Generating function |
| Use TopFD feature file | True |

**Table S6**. Parameter settings for TopCluster in TopLib

| Function | Parameters | Settings |
| --- | --- | --- |
| Precursor mass-based clustering | Clustering method | Hierarchical clustering |
|  | Distance function | Manhattan distance |
|  | Linkage method | Complete |
|  | Distance cutoff | 2.2 Da |
| MS/MS spectral clustering | Clustering method | Hierarchical clustering |
|  | Distance function | Cosine distance |
|  | Linkage method | Average |
|  | Distance cutoff value | 0.7 |
